# DeepVASP-S: a deep-learning tool for explaining steric mechanism that controls binding specificity

**DOI:** 10.1101/2023.11.04.565624

**Authors:** Yangying Liu, Houliang Zhou, Brian Y. Chen

**Affiliations:** Computer Science and Engineering Department, Lehigh University, Bethlehem, PA 18015, USA

## Abstract

DeepVASP-S is a computational tool that leverages the capabilities of convolutional neural networks (CNNs) to analyze steric aspects of protein-ligand interactions and predict amino acid contributions to binding specificity. This tool combines structural bioinformatics with machine learning to address the complex problem of understanding how specific amino acids contribute to the specificity of binding. Here, we use this tool to predict subclasses for Enolase and Serine Protease according to their binding specificity, and explain it through the underlying steric mechanism. The strength of DeepVASP-S lies in its ability to identify and highlight these specific regions, providing researchers with insights into the molecular determinants of protein function and interaction. Such information is extremely valuable for the field of drug design, as it enables the creation of more targeted and effective therapeutics with minimized side effects. The approach taken by DeepVASP-S represents a significant step forward in computational biochemistry, merging high-resolution 3D structural data with the predictive power of machine learning to unlock a deeper understanding of protein-ligand interactions.

## 1 Introduction

Members of a protein family are the product of divergent evolution caused by distinct evolutionary forces, they are homologous and tend to have similar structural characteristics. However, proteins do not work independently in living organisms, they tend to bind to other ligands (such as metal ions, nucleic acids, and inorganic or organic small molecules) to create specific interactions to achieve corresponding functions. For homology-related proteins, Even though they have similar structural characteristics, their ligands can vary [1]. Therefore homologous proteins don’t necessarily have similar ligand specificity. This indicates that structural binding-site differences are better related to ligand preferences among members than global differences. The structural binding-site specificity might be due to a small minority of binding-site residues that are specificity-determining. Because intermolecular interactions between proteins and ligands occur via amino acid residues at specific positions in the protein, usually located in pocket-like regions. Identifying those specificity-determining amino acids and studying underlying biochemical mechanisms through which they exert their effects, is crucial for understanding the evolution of ligand binding sites within a family, explaining genetic variations influence binding specificity and pathogenicity, and how binding preference might be changed through protein design.

Previous studies have made some progress in the field of ligand-binding site predictions and key amino acid identification. These methods are normally based on sequence information or structural information, employing geometry or energy feature searching, and sequence or structure similarity comparison to explore ligand binding specificity. Interactions between specific amino acids of binding sites and ligands provide underlying mechanisms for specific ligand recognition, but identifying mechanisms needs another step with human expertise. By knowing the mechanisms by which amino acids contribute to binding specificity, we can understand how genetic variations affect binding specificity and how to design experiments to efficiently explore specific drugs for the disease. If an algorithm can generate an explanation of specificity mechanisms, it could offer insights at the appropriate scale and accelerates the drug design process. Explain steric hindrance

To explain the biochemical mechanisms of key amino acids underlying selective binding, this paper proposes one method which examines only the steric hindrance of key amino acids influencing binding specificity. As one method within the Analytic Ensemble approach, this method only focuses on steric hindrance, any amino acid identified by this narrow method can be associated with a steric influence on binding specificity because the method examines no other mechanism. We begin with a training family of homology-related proteins that perform the same biochemical function, with subfamilies that prefer to act on similar but non-identical ligands.

This paper introduces DeepVASP-S, an innovative deep-learning algorithm that aims to identify the mechanisms of steric hindrance through which amino acids influence specificity in ligand binding. The algorithm achieves this objective by utilizing a solid voxel representation to effectively capture the characteristics of binding sites. DeepVASPS performs two primary functions when presented with the binding ligand of a query protein with unknown binding preferences. Firstly, it employs a three-dimensional convolutional neural network (3D-CNN) to classify the query protein into one of the subfamilies based solely on the geometric similarity of their binding sites. This classification step provides insights into the potential binding preferences of the query protein. Secondly, the algorithm leverages the gradient-weighted class activation mapping (Grad-CAM++) technique [2] to identify specific regions within the query protein that significantly contribute to its classification into a particular subfamily. By utilizing Grad-CAM++, DeepVASP-S can identify critical regions of the protein that play a vital role in its classification, potentially highlighting zones of steric crucial for selective binding. The accurate detection and definition of binding sites are fundamental to determining binding-site specificity. DeepVASP-S addresses this challenge by utilizing a solid voxel representation, which effectively represents the ligand binding sites and allows for subsequent analysis using powerful deep learning techniques.

## 2 MATERIALS AND METHODS

Proteins from the same subfamily tend to have the same binding preference but exhibit binding specificity among different subfamilies. This binding specificity can manifest in binding cavity shape, proteins from the same protein subfamily have similar binding shapes, but there are subtle differences in the binding shapes of proteins from different protein subfamilies. By leveraging this principle, we can classify proteins into correct subfamilies by contouring and representing the binding cavity shapes of different proteins. So firstly, in order to get the binding cavities shapes, we use solid volumetric representation to describe them. Then we use a deep learning method called DeepVASP-S, which employs a multi-layer CNN to classify homology-related proteins within a family into different subfamilies by recognizing their binding cavity shapes. Besides, by recognizing cavity shapes, we can not only acquire information about different proteins’ binding specificity but also get the underlying steric hindrance mechanism responsible for the specific binding preference. Furthermore, by using GradCAM++, which is a visual explanation method of CNN model predictions, we identify amino acids that exert the underlying steric hindrance mechanism to influence binding specificity to provide insight into how key amino acids affect binding affinity through steric hindrance.

### 2.0.1 Dataset construction

We selected proteins from the Enolase superfamily and Serine Proteases superfamily to construct our dataset(Table 1). For each superfamily, we included three subfamilies with different binding preferences. From the serine proteases, we selected 16 proteins from the trypsin, chymotrypsin, and elastase subfamilies. From the enolase superfamily, we selected 19 proteins from the enolase, mandelate racemase, and muconate lactonizing enzyme families. Proteins within a superfamily are sequentially nonredundant, They have a sequence identity of less than 90% with each other.

**Table 1:** PDB codes of selected families used in this study.

| Superfamily | Subfamily | PDB |
| --- | --- | --- |
| Serine Proteases | Trypsins | 2f91, 1fn8, 2eek, 1h4w, 1bzx, 1aq7, 1ane, 1aks, 1trn, 1a0j |
|  | Chymotrypsins | 1eq9, 4cha, 1kdq, 8gch |
|  | Elastases | 1elt, 1b0e |
| Enolase | Enolases | 1e9i, 1pdy, 1iyx, 1ebh, 1te6, 1w6t, 2xsx, 2pa6, 3otr, 3qn3 |
|  | Mandelate Racemases | 2ox4, 1mdr, 2og9 |
|  | Muconate Lactonizing Enzyme | 2pgw, 1bkh, 3dgb, 3i4k, 2zad, 1jpm |

**Table 2:** Prediction performance of Logistics Regression and Deep VASP S.

|  |  | PCA+LR |  | Deep VASP S |  |
| --- | --- | --- | --- | --- | --- |
|  |  | Accuracy | F1 | Accuracy | F1 |
| Enolase | Overall | 60.62418947 | 33.35703684 | 83.82322105 | 83.99165263 |
|  | Mandelate Racemase | 0 | 0 | 0 | 0 |
|  | Muconate Lactonizing Enzyme | 30.31209474 | 31.17067173 | 98.77353333 | 99.3069 |
|  | Enolase | 84.44198 | 46.4108 | 100 | 100 |
| Serine Protease | Overall | 60.62418947 | 72.58758412 | 83.976675 | 84.8998375 |
|  | Trypsin | 86.17998 | 68.16722 | 98.02344 | 98.15881 |
|  | Chymotrypsin | 39.119725 | 19.992475 | 50 | 50 |
|  | Elastase | 0 | 0 | 81.6962 | 88.40465 |

Serine protease superfamilies can have endo- or exopeptidase activity, and some of them are involved in the breakdown of proteins in the digestive system. For example, trypsins, chymotrypsins, and elastases are endoproteases that can break down polypeptides into shorter chains. However, the specificity of these three types of Serine protease is related to their substrate instead of their structural or mechanistic features [3]. Besides, for trypsins, chymotrypsins, and elastases, their substrate recognition is only related to the S1 site, while for other serine proteases, substrate recognition can extend beyond the S1 site. Substrates’ preference for these three types of serine proteases are distinct, which leads to their binding specificity. The specificity of chymotrypsin correlates with the hydrophobicity of the P1 residue, creating a large hydrophobic pocket in chymotrypsin that accounts for this specificity. Trypsin’s specificity for substrates is related to the negatively charged S1 site, therefore prefers positively charged P1 residues. Elastase prefers substrates with small aliphatic residues at P1; the S1 site of elastase is smaller than the S1 sites of chymotrypsin and trypsin [4].

Proteins in enolase superfamilies share a common partial reaction, they mediate a reaction that removes the *α*-proton from a carboxylate substrate, resulting in the creation of an enolate anion intermediate that is stabilized by binding to a crucial Mg2+ ion. Depending on the specific active site structure, these intermediates are then directed to produce different end products [5, 6]. The shared common active site is inside of a modified (b/a)8-barrel, and conserved residues are arranged around it. A second domain caps the active site, which provides substrate specificity for some enolase superfamily enzymes. Besides, the metal-coordinating residues are highly conserved, but the identity and position of the general base that abstracts the proton are specific for proteins from different subfamilies. [6].

### 2.1 Generating 3D volumetric representations for cavities

The training dataset includes proteins from a protein superfamily T that belong to different subfamilies T0,T1, ⋯,Tn, they exhibit distinct binding preferences and a known ligand-binding site, which are used for protein subclass classification. To get the binding cavity of the proteins, we firstly get 1000 conformational samples through molecular dynamic simulation, and then we abstract and represent the binding cavity shape by using genSurf.

#### 2.1.1 Conformational Sampling

Before we get the cavity samples, we need to get the conformational samples, for each protein structure in the data set, we used GROMACS 2021.4 version with cuda-11.6.0 support. We get protein structures from the PDB website, before we start the simulation, we remove all water from the structures. Firstly, we created a cubic water box and centered the protein molecule in the box with at least 1.0 nm from the box edge (-d 1.0). GROMOS96 54a7 force field is selected for generating topology. The solvent in the water box was using SPC/E, an equilibrated 3-point water model [7]. Fully periodic boundary conditions were used throughout the equilibration and simulation steps. Secondly, to make the system in the water box charge-balanced, sodium and potassium ions were then added to the solvent at a low concentration (*<* 0.1% salinity) to neutralize the net charge on the protein. Thirdly, energy minimization using a steepest descent algorithm is then performed for the entire system. Isothermal-Isobaric (NPT) equilibration is performed in four 250 picosecond steps to allow the solvent to equilibrate temperature and pressure prior to the primary simulation. Starting at 1000 kJ/(mo nm), each step reduced the position restraint force by 250 kJ/(mol nm) over the 1 nanosecond minimization period. Backbone position restraints were released for the primary NPT simulation. System energies were generated at the start of the equilibration phase. The initial pressure was 1 bar and the Parrinello-Rahman algorithm was used for pressure coupling. The initial temperature was 300 Kelvin and the Nos’e-Hoover thermostat was used for temperature coupling. All temperature and pressure scaling was performed isotropically. The P-LINCS bond constraint algorithm was used to update [8]. Electrostatic interaction energies were calculated by particle mesh Ewald summation (PME) [9]. Full MD simulation is started using the atomic positions and velocities of the final equilibration state. The total simulated duration of the molecular dynamics simulation was 100 nanoseconds, with 1 femtosecond steps. OpenMPI was used for node and inter-process communication. After simulations were completed, the trajectory file was converted to a simple Protein Data Bank format with atom positions only. The waterbox was removed and at each timestep, the protein was rigidly superposed to the original orientation. From these timesteps, we selected 1000 conformational samples.

#### 2.1.2 Structural Alignment and Binding Cavity Representation

After conformational sampling, we used resAlign to align structures to pivot protein. For serine proteases, bovine chymotrypsin (with PDB code 8gch) was the pivot structure, and for enolases, the pseudomonas putida mandelate racemase (PDB code 1mdr) was the pivot structure. These particular reference proteins, already crystallized with ligands, were selected to localize a binding site. After aligning with the pivot structure, for each protein, its 1000 confirmational samples are aligned with their corresponding aligned structure.

Then, the genSurf tool was employed to generate a 3D solid representation for the binding cavity. Firstly, we generated spheres with 5.0 Åin radius, centered around the atoms of the ligand, CSG union of all spheres is used to form a ligand representation as “U”. In parallel, the molecular surface (“S”) and the envelope surface (“E”) of the protein conformation were determined by applying the classic rolling probe algorithm. Here, “S” and “E” surfaces were defined by probes of 1.4 Åand 5.0 Åradius, respectively, E represents the region inside the protein, including the cavity. Using CSG, we compute (*U* − *S*) ∩ *E* to produce the cavity [10].

#### 2.1.3 Voxelized Binding Site Representations

After the cavities within the protein structures are determined, they are converted into a grid-based representation for 3D Convolutional Neural Network (3D-CNN) analysis. Generating a bounding box that is consistent across all proteins and can accommodate all proteins from the same superfamilies within it. This bounding box is then segmented into a grid of small, equally-sized cubes, each with a volume of 0.5Åper side. To accommodate a whole number of these cubes, the bounding box is adjusted with slight padding as necessary. By performing CSG operations with the cubes, we can calculate the portion of the cavity’s volume that occupies each voxel. The resulting array of voxel volumes, a tensor, is then used as input data for training or classifying with the 3D-CNN.

#### 2.1.4 Generated frequent regions for cavities

A “frequent region” is characterized as a spatial zone that is solvent-accessible in more than k out of N conformational samples. In this definition, k is the overlap threshold, which is a predetermined input value, and N represents the total number of samples. These frequent regions, therefore, pinpoint cavities that are solvent-accessible in a meaningful number of conformations, avoiding those regions that are only occasionally exposed due to rare structural variations. Here, we set k as 15 and randomly select 15 cavity representations from 1000 samples to get the intersection of 15 cavities, then repeat the random selection and intersection generation process 500 times to get 500 intersections, and then we get the union of 500 intersections to produce frequent region. In total, we got 500 frequent region representations for 3D-CNN analysis.

### 2.2 CNN structure

The 3D-CNN model within our DeepVASP-S framework is tailored to process voxel-based inputs. A key consideration in shaping the DeepVASP-S CNN structure is the high resolution of the input cavies, which span 30 to 42 cubes across each spatial dimension. This results in a substantially high neuron count in each layer, a figure that surpasses what’s typically encountered in standard 2D image processing. In attempts to manage this complexity, reducing neuron numbers through increased voxel size was tested, but such reductions were found to adversely affect the model’s classification precision. Thus, to maintain high accuracy, the model retains full input resolution and adopts a comparatively less deep architecture, akin to that of the classic networks it emulates. The network culminates in a dense layer utilizing a softmax activation to categorize the inputs into one of three subfamily classifications, aligning with the divisions within our training dataset.

### 2.3 GradCAM++ class-activation map

Grad-CAM++ (Gradient-weighted Class Activation Mapping) is an advanced visualization technique used in the field of machine learning, particularly in the context of Convolutional Neural Networks (CNNs). It is an extension of the original Grad-CAM method and provides an improved way to visualize and understand which parts of a given input image are most important for the predictions made by a CNN. It provides better visual explanations of CNN model predictions.A weighted combination of the positive partial derivatives of the last convolutional layer feature maps with respect to a specific class score as weights are used to generate a visual explanation for the corresponding class label. Here, we try to find the most important regions and amino acids that contribute to the classification.

## 3 RESULTS

### 3.1 Classification

We evaluated the performance of DeepVASP-S for predicting the specificity category of each member of both training families. The average accuracy and F1-score of the compared methods are presented in Table **??**.

Overall, DeepVASP-S clearly outperformed PCA with logistics regression on three out of four datasets and performed worse on the fourth dataset. These findings point to lower consistency in the classification performance of PCA relative to DeepVASP-S. Compared to PCA-based logistic regression, another benefit of the proposed DeepVASP-S model is that it maintains the adjacency structure of the voxel data, rather than vectorizing it, to support explainability. We exploit this advantage using Grad-CAM++ below.

### 3.2 Mechanism

We hypothesize that the most salient voxels identified by DeepVASP-S will be regions of specificity of the protein caused by the steric hindrance mechanism. Thus, we also evaluated the accuracy of the most salient voxels and verified them against experimentally established findings in the literature. Overall, we observed that the most salient voxels appear near charged amino acids that are known to affect specificity. The clearest example of this observation is the case of Val216 in trypsin. This amino acid has been experimentally established to play a pivotal role in binding specificity. The most salient region for classification is a region near Val216 (Fig. 1). It is clear that the most salient voxels are identifying a region that enables specificity in Elaseteas.Val216 is an amino acid exited in trypsin instead of elastase, it blocks the saliency voxels of elastase and therefore they have different binding preferences. It is clear that this region plays a crucial role in distinguishing the subfamilies of the serine proteases, and that saliency mapping is able to detect steric influences on specificity.

**Figure 1:**
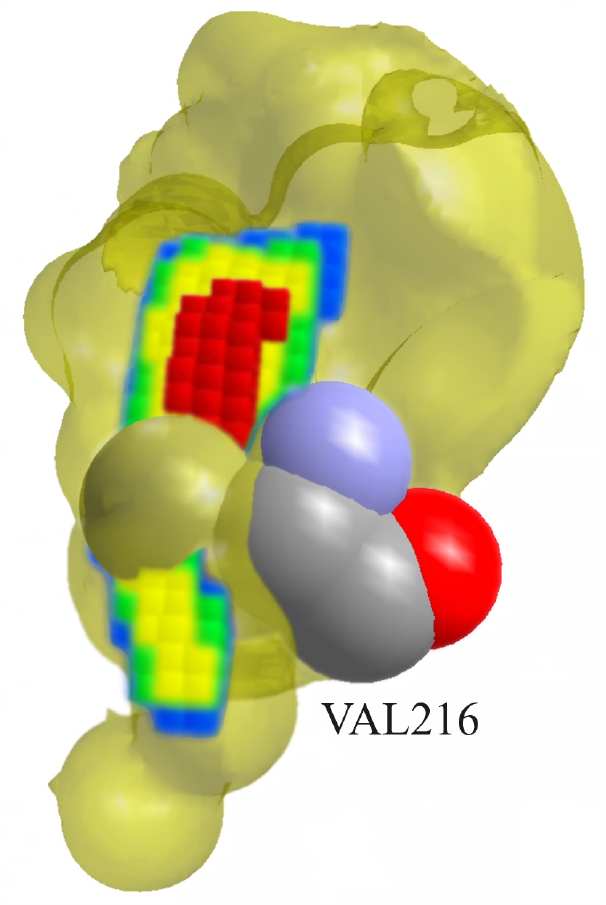
Saliency Map of Elastase 1ane with VAL216 of Trypsin 1b0e

## 4 CONCLUSION

DeepVASP-S is an efficient computational tool that use convolutional neural networks (CNNs) to analyze steric aspects of protein-ligand interactions and predict amino acid contributions to binding specificity. This tool combines structural bioinformatics with machine learning to address the complex problem of understanding how specific amino acids contribute to the specificity of binding, which provides an accurate classification and generates a meaningful explanation for binding specificity in steric aspect.

## Notes

### Competing Interest Statement

The authors have declared no competing interest.

